# Expansion load: recessive mutations and the role of standing genetic variation

**DOI:** 10.1101/011593

**Authors:** Stephan Peischl, Laurent Excoffier

## Abstract

Expanding populations incur a mutation burden – the so-called expansion load. Previous studies of expansion load have focused on co-dominant mutations. An important consequence of this assumption is that expansion load stems exclusively from the accumulation of new mutations occurring in individuals living at the wave front. Using individual-based simulations we study here the dynamics of standing genetic variation at the front of expansions, and its consequences on mean fitness if mutations are recessive. We find that deleterious genetic diversity is quickly lost at the front of the expansion, but the loss of deleterious mutations at some loci is compensated by an increase of their frequencies at other loci. The frequency of deleterious homozygotes therefore increases along the expansion axis whereas the average number of deleterious mutations per individual remains nearly constant across the species range. This reveals two important differences to co-dominant models: (i) mean fitness at the front of the expansion drops much faster if mutations are recessive, and (ii) mutation load can increase during the expansion even if the total number of deleterious mutations per individual remains constant. We use our model to make predictions about the shape of the site frequency spectrum at the front of range expansion, and about correlations between heterozygosity and fitness in different parts of the species range. Importantly, these predictions provide opportunities to empirically validate our theoretical results. We discuss our findings in the light of recent results on the distribution of deleterious genetic variation across human populations, and link them to empirical results on the correlation of heterozygosity and fitness found in many natural range expansions.

## Introduction

Identifying and understanding the ecological and evolutionary processes that cause range expansions, range shifts, or contractions has a long tradition in evolutionary biology (Darwin 1859; MacArthur 1972; Sexton *et al.* 2009). More recently, the growing appreciation of the consequences of dynamic range margins on the ecology, population genetics, and behavior of species has changed our views about several evolutionary processes, such as the evolution of dispersal (Phillips *et al.* 2006; Shine *et al.* 2011; Lindström *et al.* 2013), life-history traits (Phillips *et al.* 2010), and species range limits (Peischl *et al.* 2014).

The evolutionary processes at the margins of expanding populations allow neutral genetic variants to quickly spread into new territories (Klopfstein *et al.* 2006), a phenomenon called “gene surfing”. Gene-surfing of neutral variation has been investigated both theoretically (Hallatschek and Nelson 2008; Excoffier *et al.* 2009; Slatkin and Excoffier 2012) and empirically (Hallatschek and Nelson 2008; Moreau *et al.* 2011; Graciá *et al.* 2013). Gene surfing can also affect the spread of selected variants (Travis *et al.* 2007; Burton and Travis 2008; Lehe *et al.* 2012; Peischl *et al.* 2013; Peischl *et al.* 2014). Population-genetics models of range expansions predict that expanding populations incur a mutation burden – the “expansion load” (Peischl *et al.* 2013). Expansion load is a transient phenomenon, but it can persist for several hundreds to thousands of generations, and may limit the ability of a species to colonize new habitats (Peischl *et al.* 2014).

Previous studies of expansion load assumed that mutations were co-dominant. An important consequence of this assumption is that standing genetic variation has no effect on the dynamics of mean fitness at the front of expanding populations (Peischl *et al.* 2013). In particular, the total number of mutations per individual, and hence the individual’s fitness, remains approximately constant if new mutations are ignored (Peischl *et al.* 2013; Peischl *et al.* 2014). In additive models, expansion load thus stems exclusively from the accumulation of new mutations that occur in individuals living at the front of the expansion.

Empirical evidence for expansion load may come from humans, where a proportional excess of deleterious mutations in non-African populations has been found (Lohmueller *et al.* 2008; Subramanian 2012; Torkamani *et al.* 2012; Peischl *et al.* 2013; Fu *et al.* 2014; Lohmueller 2014). Importantly, when focusing on mutations that occurred during or after the out-of Africa expansion, the excess of deleterious variants is not restricted to rare variants (Peischl *et al.* 2013). This suggests that proportionally more deleterious mutations have risen to high frequencies in human populations located in newly settled habitats. In contrast to what would be expected from expansion-load theory, a recent analysis found no significant differences in the average allele frequency of predicted deleterious alleles (Simons *et al.* 2014). The average number of predicted deleterious mutations carried by an individual is, however, significantly larger in non-Africans (Fu *et al.* 2014). In addition, non-African individuals have significantly more loci homozygous for predicted deleterious alleles than African individuals (Lohmueller *et al.* 2008; Subramanian 2012; Fu *et al.* 2014; Lohmueller 2014). The debate whether human past demography affected the efficacy of selection and the spatial distribution of mutation load is thus still ongoing (Lohmueller 2014).

There is mounting evidence that deleterious mutations tend to be recessive (Agrawal and Whitlock 2011). Importantly, if mutations are completely recessive the number of deleterious mutations per individual is not informative about the mutation load (Kimura *et al.* 1963). Instead, mutation load is determined by sites that are homozygous for deleterious alleles. Thus, if mutations are (partially) recessive, the genotypic composition of deleterious genetic variation is more important than the total number of deleterious mutations carried by an individual. Range expansions are known to affect the genotypic composition of neutral standing genetic variation (Excoffier *et al.* 2009). The role of standing genetic variation in models of expansion load remains, however, unclear if mutations are recessive.

We investigate here the effect of recessive mutations on the dynamics of expansion load. In particular, we use individual based-simulations to investigate the role of standing genetic variation, the width of the habitat, and the composition of expansion load with respect to allele frequencies and mutational effects.

## Model and Results

### Model

We model a population of diploid monoecious individuals that occupy discrete demes located on a one-or two-dimensional grid (Kimura and Weiss 1964). Generations are discrete and non-overlapping, and mating within each deme is random. Mating pairs are formed by randomly drawing individuals (with replacement) according to their relative fitness, and each mating pair produces a single offspring. The process is repeated *N* times, where *N* is the total number of offsprings of the parental generation, leading to approximately Poisson-distributed numbers of offspring per individual. Individuals then migrate to adjacent demes with probability *m* per generation. Migration is homogeneous and isotropic, except that the boundaries of the habitat are reflecting, i.e., individuals cannot migrate out of the habitat.

Population size grows logistically within demes. The expected number of offspring in the next generation produced by the *N_j_* adults in deme *j* is

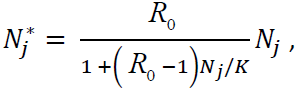

where *R*_0_ is the fundamental (geometric) growth rate and *K* is the deme’s carrying capacity (Beverton and Holt 1957). To model demographic stochasticity, the actual number of offspring, 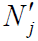, is then drawn from a Poisson distribution with mean 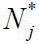.

The relative fitness of individuals is determined by *n* independently segregating biallelic loci. The alleles at locus *i* are denoted *a*_*i*_ (wildtype) and *A*_*i*_(derived). Mutations occur in both directions and the genome wide mutation rate is *u*; in each new gamete *k* randomly chosen sites change their allelic state, where *k* is drawn from a Poisson-distribution with mean *u*. The fitness contributions of the genotypes *a*_*i*_*a*_*i*_, *a*_*i*_*A*_*i*_ and *A*_*i*_*A*_*i*_ at locus *i* are 1, 1 − *hs*_*i*_, and 1−*s*_*i*_, respectively. Here *s*_*i*_ denotes the strength of selection at locus *i* and *h* is the dominance coefficient. Fitness effects are multiplicative across loci, such that the fitness of an individual is given by *w*=∏_*i*_*w*_*i*_, where *w*_*i*_ is the fitness effect of the *i*th locus of the focus individual, i.e., there is no epistasis. In the following we will focus on co-dominant (*h* = 0.5) or recessive (*h* = 0) mutations. We assume that mutation effects are drawn from the same distribution of fitness effects for all individuals (independently from their current fitness).

We perform individual-based simulations of the above described model in 1D or 2D habitats. Our simulations start from ancestral populations located in 10 leftmost (rows of) demes of the range. After a burn-in phase that ensures that the ancestral populations are at mutation-selection-drift balance, the population expands from left to right until the habitat is filled. Because we are mainly interested in the role of standing genetic variation, we focus on relatively short expansions, i.e., colonization of a 1x50 (1D) or a 20x50 (2D) deme habitat. The long-term dynamics of expansion load have been studied elsewhere (Peischl *et al.* 2013; Peischl *et al.* 2014).

### Impact of standing genetic variation on expansion load

For simplicity, we first consider expansions along a one-dimensional habitat and assume that all mutations have the same effect, i.e., we set *s*_*i*_ = *s*. If mutations are co-dominant (*h* = 0.5), expansion load is caused exclusively by the establishment of new mutations occurring during the expansion, and standing genetic variation has a negligible effect on the dynamics of mean fitness (Peischl *et al.* 2013). Mean fitness at the wave front decreases at a constant rate over time (Figure 1), and the rate at which mean fitness decreases per generation is proportional to the number of new mutations entering the population per generation (Peischl *et al.* 2013).

**Figure 1:**
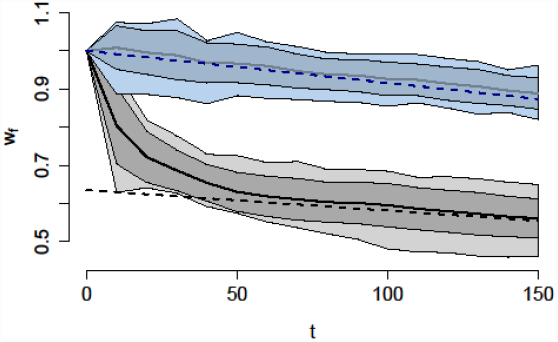
Evolution of mean fitness at the wave front. Dashed lines show analytical prediction for the evolution of the mean fitness due to de-novo mutations (see Peischl et al. 2013). Simulations show results for the combination of standing and new genetic variation. Gray shaded areas and black lines show results for recessive mutations (*h* = 0), and blue shaded areas and lines show results for additive co-dominant mutations (*h* = 0.5). Solid lines indicate the average mean fitness from 50 simulations, and dark and light shaded areas indicate ± one standard deviation and the minimum and maximum of mean fitness, respectively. Other parameter values are *n* = 1000, *K* = 100, *u* = 0.1, *m* = 0.1, *s* = 0.01, *R* = 2.

The dynamics of expansion load changes dramatically if mutations are recessive (Figure 1). The analytical approximation obtained in Peischl *et al.* (2013), which ignores standing genetic variation, is a poor fit to the observed dynamics of mean fitness (Figure 1). In the first few generations mean fitness decreases much faster than predicted by analytical theory for the accumulation of new mutations (cf. solid and dashed black lines in Figure 1). The rate at which expansion load is created then slows down and gradually approaches the analytical prediction. Then, changes in expected mean fitness arise exclusively from new mutations (cf. solid and dashed black lines for *t* > 50 in Figure 1). This shows that standing genetic variation plays an important role in the establishment of expansion load if mutations are recessive, especially during early phases of expansions.

We next investigate the evolution of the genotypic composition of standing genetic variation on the expansion front. In general, we find that the average number of heterozygous loci per individual decreases during the expansion, whereas the number of loci that are homozygous for the derived allele increases (Figure 2). Because we simulated a fixed number of loci, the derived allele frequency shown in Figure 2 is proportional to the average number of mutations carried by an individual. Thus, Figure 2 shows that the total number of mutations per individual remains nearly constant during the expansion (Figure 2). Strong genetic drift is therefore the major force driving the evolution of genotype frequencies at the wave front. At any given locus, mutations are either lost or fixed over the course of the expansion, and the probability of fixation of a given mutation is close to its initial frequency (Peischl *et al.* 2013), suggesting that deleterious mutations are behaving like neutral mutations on the wave front. In 2D expansions, the dynamics of genotype frequencies are qualitatively very similar to 1D expansions (Figure S1).

**Figure 2:**
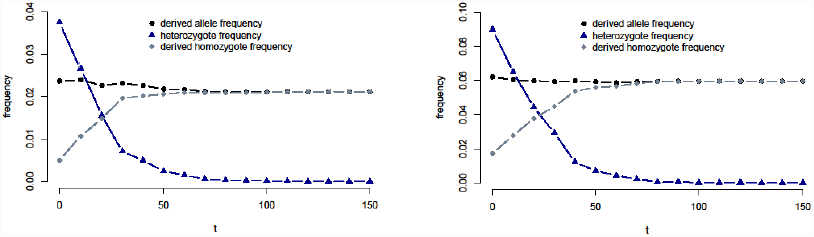

Evolution of standing genetic variation on the wave front. Panel A shows results for co-dominant mutations and panel B for recessive mutations. Parameter values are as in Figure 1.

The nearly neutral evolution of allele frequencies on the expansion front reveals a critical role of the degree of dominance on the build-up of the expansion load. If mutations are co-dominant, the fitness of an individual is determined by the total number of mutations it carries (Wright 1930). Thus, Figure 2 shows that standing genetic variation has a negligible impact on fitness (Figure 2A). In contrast, if mutations are recessive, the fitness of an individual is determined by its number of loci homozygous for the derived allele. Because the number of derived homozygous loci per individual rapidly increases at the front of the expansions, standing genetic variation has a severe effect on fitness if mutations are recessive (Figures 1 and 2B).

## Gene flow on the wave front of 2D expansions restores diversity and fitness

In the following section, we focus on completely recessive mutations (*h* = 0). Figure 3 shows an example of the evolution of the mean fitness during an expansion in a 2D habitat (20x50 demes). As in 1D expansions, the mean fitness drops to low levels on the expansion front within the first few (≈30) generations and then continues to gradually decreases at a slower rate. There is however a considerable variation in fitness across the wave front of 2D expansions (fitness-differences of more than 40%, Figures 3 and 4). At the end of the expansion (Figure 3, *T* =150), we find a high-fitness ridge along the expansion axis in the central part of the newly settled species range, surrounded by sectors of low fitness on the lateral edges of the species range. This is partially caused by the lack of immigrants at the lateral edge of the species range (boundary effect). However, the location of the high-fitness ridge varies across simulation runs, suggesting that a boundary effect alone cannot explain the observed patterns (Figure S2).

**Figure 3:**
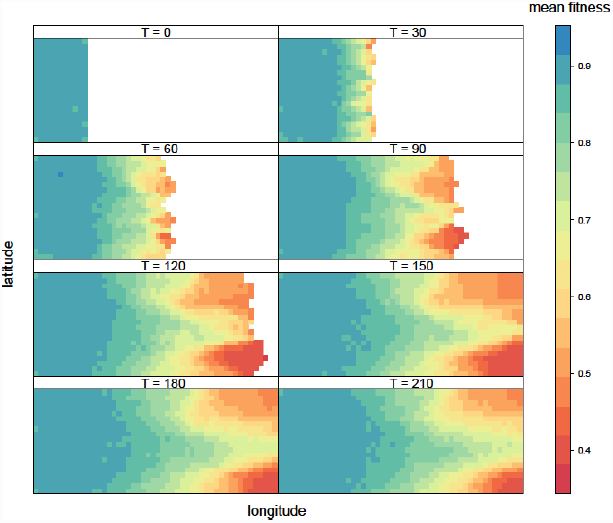
Evolution of mean fitness during range expansion. The simulated grid is 20x50 demes. Mutations are recessive and parameter values are as in Figure 1.

**Figure 4:**
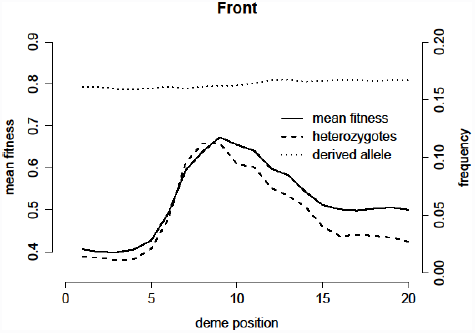

Genetic properties of the demes located on the wave. The figure shows the mean fitness, the heterozygosity, and the derived allele frequency at the front of the expansion when the habitat has just been fully colonized. The deme mean fitness on the wave front correlates with heterozygosity, but not with derived allele frequency. Statistics were computed from the simulation shown in Figure 3 at generation 150.

Figure 4 shows the variation in fitness, heterozygosity, and derived allele frequency across the wave front at the end of the expansion shown in Figure 3. We find that the average number of mutations per individual is uniform across the expansion front, which means that the variation in fitness across the expansion front is not driven by a differential accumulation of mutations. Heterozygosity, on the other hand, correlates strongly with mean fitness (cf. solid and dashed line in Figure 4). This observation suggests that different mutations establish in different parts of the wave front, and that gene flow between demes restores heterozygosity, which masks the effect of deleterious recessive mutations. These results show that heterozygosity-fitness correlations (HFC) are readily created during range expansions. We indeed find a strongly positive HFC at the front of the expansion (Figure 5A, *R*^2^=0.526., *slope = 0.32*, *p*<10^−16^), but not in the ancestral population(Figure 5B, *R*^2^=0.001, *slope = 0.03*, *p* >0.07). Interestingly, weaker but similar correlations are found at the individual level within demes (mean slope ≈0.1, *p* <0.05 in 81% of all simulated demes).

**Figure 5:**
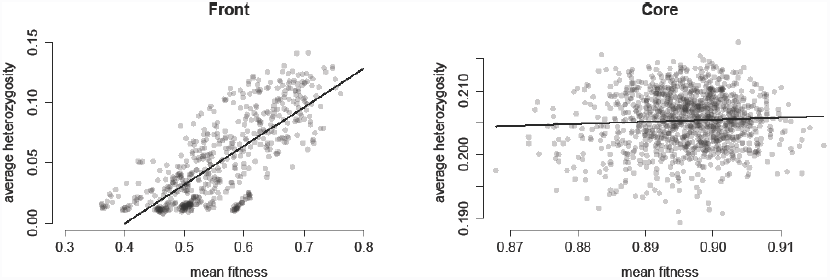

Heterozygosity-Fitness correlations (HFC). (A) HFC on the expansion front at generation *T* =120 (*R*^2^ =0.526, *slope = 0.32*, *p* <10^−16^). (B) No significant HFC in core populations before the onset of the expansion (*R*^2^ =0.001, slope = 0.03, *p* >0.07). Each point represents a single deme. The results from 10 simulation replicates are shown. Parameter values are as in Figure 3

## Expansion load is driven by a few mutations occurring at high frequency

So far we assumed that all mutations had the same effect *s*. To investigate the composition of expansion load with respect to mutation fitness effects, we now consider the case where mutation fitness effects are drawn from an exponential distribution with mean *s*. Figure 6A and B show the site frequency spectrum (SFS) observed in core and front populations, respectively. In core populations, the SFS shows the pattern expected for sites under negative selection (Bustamante *et al.* 2001), with a large excess of low frequency variants. On the wave front, the total number of segregating sites is reduced in marginal populations (cf. Figures 6 A and B). More interestingly, we see a markedly different SFS, with, as compared to neutral expectation, a clear deficit of rare and intermediate frequency variants and an increase in high frequency variants (Figure 6B). Thus, even though fewer polymorphic sites with deleterious variants are found in more recently colonized areas than in the ancestral region, the alleles at polymorphic sites tend to be at higher frequency in more recently colonized populations.

**Figure 6:**
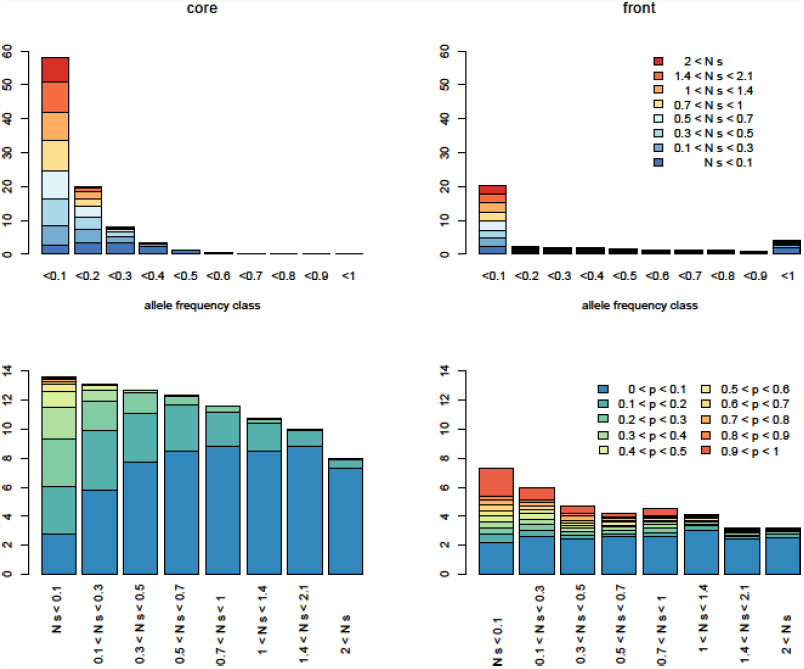

Distribution of average number of polymorphic loci per individual. The distribution is stratified for allele frequencies (top row) and mutation effects (bottom row). Results were recorded 150 generations after the onset of the expansion, which is shortly after the habitat was colonized completely (mean time to colonization ≈130 generations, see also Figure 3). Panels (A) and (C) show results for a core population (coordinates 10, 5), (B) and (D) front population (coordinates 10,45). Mutations are recessive and their effects are drawn from an exponential distribution with mean s = 0.01. Other parameter values are as in Figure 1.

Figure 6 C and D show the distribution of polymorphic loci stratified according to their mutation effect sizes. The eight mutation effect classes have been defined such that they represent the 8-quantiles of the DFE, i.e., the rate at which mutations of a given category enter the population are equal for all categories. As expected, we find that the number of polymorphic loci generally decreases with increasing mutation effect size, and that large effect mutations tend to be present at lower frequencies than low effect mutations (Figure 6 C and D). Compared to core populations, the allele frequencies at polymorphic sites on the wave front tend to be larger across all mutational effect categories. Furthermore, the increase in allele frequency is most pronounced for small effect mutations. Thus, expansion load is driven mainly by mildly and moderately standing deleterious mutations (i.e., up to *Ns* <2 for the parameter values used in Figure 6) that rise to high frequency during the expansion.

## Discussion

We have investigated here the dynamics of an expansion load caused by recessive mutations. Using individual-based simulations we have shown that shifts in the genotypic composition of standing genetic variation can lead to a rapid drop of mean fitness at the onset of an expansion (see Figures 1 and 2) without necessarily affecting the total number of deleterious alleles per individuals (see Figure 2). The total expansion load resulting from standing genetic variation is limited by the initial frequency of deleterious mutations (see Figure 2). Thus, if many loci are polymorphic for deleterious variants at the onset of the expansion, the (recessive) expansion load from standing genetic variation can be the dominating the total mutation load (see Figure 1). Even though these results have been inferred by assuming that all deleterious mutations were recessive, we would predict that a similar phenomenon, though of lesser amplitude, would occur if only some of the mutations would be fully or partially recessive.

The effect of range expansions on deleterious genetic diversity is also reflected in the site frequency spectrum (SFS, see Figure 6). As compared to stationary populations in the core of the species range, populations from more recently colonized areas have fewer segregating sites, but proportionally more high and low frequency variants (cf. Figure 6A and B). These differences in the SFS of core and front populations should provide an opportunity to evidence expansion load from sequence data, and to infer important quantities such as the distribution of fitness effects (Keightley and Eyre-Walker 2007; Boyko *et al.* 2008; Racimo and Schraiber 2014). The development of statistical and computational methods able to infer parameters under spatially explicit models including range expansions and selection remains, however, a major challenge (Sousa *et al.* 2014).

Interestingly, human genomic data are consistent with our predictions for genomic signatures of expansion load. In particular, the number of segregating sites is higher in African populations than in non-African populations (Lohmueller *et al.* 2008), non-African populations show an excess of low-frequency and high-frequency deleterious alleles (Lohmueller *et al.* 2008; Fu *et al.* 2014), the average number of sites that are homozygous for predicted deleterious variants sites is larger in non-African individuals (Fu *et al.* 2014), and the average number of predicted deleterious mutations per individual is slightly, but significantly, larger in non-Africans (Fu *et al.* 2014). Determining mutation load (or, alternatively, fitness) from genomic variation data is, however, an intrinsically difficult problem because mutation load depends on many unknown parameters (selection coefficients that may vary over space and time, epistatic interactions, dominance relationships, etc.), and the relevance of comparing a population with deleterious mutations to a theoretical population free of such mutations is questionable (Lesecque *et al.* 2012). Testing theoretical predictions of the effect of a range expansion on functional diversity with human genomic data might nevertheless be extremely useful to substantially increase our understanding of the complex interactions of demography and selection.

We assumed here that selection was soft, i.e., demographic parameters are independent of fitness (Wallace 1975), but it would be interesting to extend our results to models of hard selection, where mutation load on the front can stop an expansion and even drive parts of the species range to extinction (Peischl *et al.* 2014). Our results suggest that admixture during range expansions, or secondary contact between expanding lineages, could mitigate expansion load and prevent marginal populations from collapsing. A previous study of range expansions under an additive model with hard selection has shown that suppressing recombination at the wave front can have beneficial effects for the spread of high fitness lineages (Peischl *et al.* 2014). Recombination modifiers, such as inversions, could have a similar effect if mutations are recessive and facilitate the spread of admixed lineages. An interesting example for studying the potentially beneficial role of admixture and suppressed recombination during range expansions is from the clam genus *Corbicula*, which includes both sexual and asexual (androgenetic diploid) lineages. Sexual populations are restricted to their native Asian areas, but the androgenetic lineages are widely distributed and extend as far as in America and Europe where they are invasive (Pigneur *et al.* 2014). Intriguingly, the invasive lineages also show an excess of heterozygosity, which is preserved through clonal reproduction. No such excess of heterozygosity is found in the native range, suggesting that the combination of asexual reproduction and high heterozygosity may have been key drivers of the invasion.

An interesting prediction of our model is that if a given proportion of deleterious mutations are recessive, then heterozygosity-fitness correlations (HFC) should naturally occur in populations that have recently expanded their range (see Figures 4 and 5A). Importantly, the positive correlation between heterozygosity and fitness in recently colonized areas can be observed at both the individual and the population level (Fig. 5). Even though our simulations modeled a single expansion in a 2D habitat, we would expect similar HFCs if there was a secondary contact between expanding populations from different areas (e.g., from different LGM refuge areas). The HFC should be even stronger in the case of a secondary contact, because the isolation between expanding lineages should be larger and different recessive alleles could have fixed in different refugia or during the expansion from these refugia. HFC have been observed in many cases of natural range expansions and invasive species (Chapman *et al.* 2009) but their underlying mechanisms and their role during range expansions and invasions are still unclear (Szulkin *et al.* 2010; Rius and Darling 2014)-A particularly interesting example of HFC is found in the invasive weed *Silene vulgaris*, where, as predicted by our model (see Figure 5), HFC correlations are observed in the recently invaded North American range, but not in their native European range. It remains however unclear whether admixture between divergent lineages has indeed a causal role in range expansions. A combination of transplantation experiments and genomic data analyses could certainly be used to test the predictions of our model.

In summary, we have investigated here the evolution of standing genetic variation during range expansions, the dynamics of mean fitness on the expansion front if mutations are recessive, and the genomic signature of range expansions. Importantly, our results make predictions that can be tested in natural populations. Empirical validation of our results would increase our understanding of the interactions of demography and selection (Lohmueller 2014), and could help us identifying key drivers of range expansions and biological invasions (Rius and Darling 2014).

## Acknowledgements

SP was supported by a Swiss NSF grant No. 31003A-143393 to LE. We thank Vitor Sousa and Isabelle Duperret for stimulating discussions on this topic.

## Supplementary Figures

**Figure S1.**
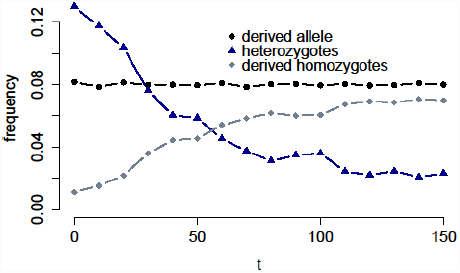
Evolution of genotype frequencies at the front of an expansion in a 2D habitat of 20x50 demes. Parameter values are as in Figure 1B.

**Figure S2.**
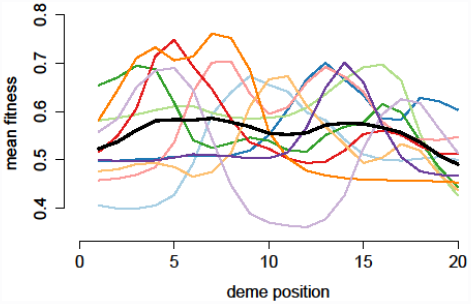
Mean fitness at the front of the expansion. The figure shows the mean fitness at the front of the expansion (at generation 150). Colored lines are the results from 10 simulation runs, solid black line shows the average over all simulation runs. Parameter values are as in Figure 3.

